# Reproducible model development in the Cardiac Electrophysiology Web Lab

**DOI:** 10.1101/257683

**Authors:** Aidan C. Daly, Michael Clerx, Kylie A. Beattie, Jonathan Cooper, David J. Gavaghan, Gary R. Mirams

**Affiliations:** Computational Biology, Department of Computer Science, University of Oxford, Oxford, UK; Research IT Services, University College London, London, UK; Centre for Mathematical Medicine & Biology, School of Mathematical Sciences, University of Nottingham, Nottingham, UK

## Abstract

The modelling of the electrophysiology of cardiac cells is one of the most mature areas of systems biology. This extended concentration of research effort brings with it new challenges, foremost among which is that of choosing which of these models is most suitable for addressing a particular scientific question. In a previous paper, we presented our initial work in developing an online resource for the characterisation and comparison of electrophysiological cell models in a wide range of experimental scenarios. In that work, we described how we had developed a novel protocol language that allowed us to separate the details of the mathematical model (the majority of cardiac cell models take the form of ordinary differential equations) from the experimental protocol being simulated. We developed a fully-open online repository (which we termed the Cardiac Electrophysiology Web Lab) which allows users to store and compare the results of applying the same experimental protocol to competing models. In the current paper we describe the most recent and planned extensions of this work, focused on supporting the process of model building from experimental data. We outline the necessary work to develop a machine-readable language to describe the process of inferring parameters from wet lab datasets, and illustrate our approach through a detailed example of fitting a model of the hERG channel using experimental data. We conclude by discussing the future challenges in making further progress in this domain towards our goal of facilitating a fully reproducible approach to the development of cardiac cell models.

## 1. Introduction

The problem of reproducibility in science is becoming well known and of great concern to researchers, funders and publishers. Computational studies are immune from many of the statistical traps (Ioannidis, 2005) that traditionally lead to problems in reproducing studies. Yet, when asked “what factors contribute to irreproducible research?” 45% of scientists replied that “methods and code being unavailable” always/often contributed, and 40% replied that “raw data availability” always/often contributed. These figures rose to 80% for each “sometimes” contributing to irreproducible research (Baker, 2016). Our field of cardiac electrophysiology modelling is not immune to these factors.

We define *reproduction* as “independent re-implementation of the essential aspects of a carefully described experiment,” which we contrast with *replication*, defined as “re-running a simulation in exactly the same way” (Howe, 2012; Cooper et al., 2015b). Aiming for replication to get precisely the same answer, in exactly the same way, by providing the code and library versions used in a study is a worthy goal in itself; it serves an important purpose in guaranteeing a minimum level of methods reporting, and may be an important stepping stone to allowing people to adapt your code for their studies. However, enabling reproduction of the process that was followed for the same *and* similar settings (without the need for code adaptation, or even making the process possible using many different codes) is a far more useful aim than replication of a single computational study (Drummond, 2009).

This paper will discuss the issue of reproducible model development, which is distinct from reproducing a computational study undertaken with a given model. Reproducing a study with a given model may assume that the model is a fixed entity (with its parameters and equations pre-defined). *Reproducible model development* means being able to recreate the process of building a model from data — fitting parameter values within equations, or even selecting the set of equations. Reproducible model development is a requirement for new science. All of the following uses of a previously published model will be performed better if we know how a model was developed: (i) using an existing model within a new simulation study/conditions, (ii) fitting an existing model to new datasets for new experimental situations (e.g. cell types, species, temperatures); and (iii) extending an existing model to explore new experimental phenomena. We will elaborate on why these applications require a well-documented and reproducible model development process below.

We focus this article on development of mathematical cardiac electrophysiology models. The first model of the cardiac action potential was created by Denis Noble (Noble, 1960, 1962) for Purkinje fibres, based on the Hodgkin and Huxley (1952) paradigm. The field of cardiac cell modelling has blossomed into one of the most popular and mature areas of systems biology research. In the ensuing decades, the scope and number of these models has increased dramatically, spurred on by a desire to recreate and understand electrophysiological phenomena across a range of species, cell types, and experimental conditions (Noble and Rudy, 2001; Noble et al., 2012). This research has been undertaken by a diverse global research community, with members of disparate institutions seeking to make improvements to existing model formulations through iterative studies in modelling and experimentation. Despite the availability of community-focused tools, we believe there remain problems with the ways modelling studies are reported that limit the usefulness, reproducibility and re-usability of our models. These problems certainly exist in many other domains, and many of the ideas in this manuscript should be directly transferable to other differential equation-based biological models. Cardiac modelling is a good area to focus on first because of its maturity, its standardisation in terms of a modular Hodgkin & Huxley-derived approach, and its importance in scientific and clinical applications.

There are at least three aspects to replicability and reproducibility in computational models that we would like to distinguish between. We have outlined these aspects in Table 1, and we discuss the entries in each row below:

1. **Models** — in order to facilitate cooperation among this physically dispersed research community, a number of tools have been developed to aid in the representation and exchange of models. Provision of software code that states equations and parameter values in an unambiguous format is an excellent and welcome step in model reporting. While providing code is a prerequisite for basic replication, as a reporting practice on its own it limits our ability to apply models to new systems without substantial alterations. So more generically and usefully, reproducibility is provided by model markup languages such as CellML/SBML (Lloyd et al., 2004; Garny et al., 2008; Hucka et al., 2003) and model repositories such as the Physiome Model Repository or BioModels Database (Yu et al., 2011; Chelliah et al., 2015) that provide public and curated reference versions of models. These repository versions of models can be used to auto-generate code in many different programming languages that can provide reproduction. Open-source software libraries such as Chaste (Mirams et al., 2013; Cooper et al., 2015a), OpenCOR (Garny and Hunter, 2015), Myokit (Clerx et al., 2016), and COPASI (Sütterlin et al., 2012) have been developed to perform simulations on models specified in these formats.
2. **Protocols** — it is helpful to maintain a separation between the notions of a ‘model’, the underlying mathematical equations thought to represent the system, and a ‘protocol’, the manner in which the system is interrogated/stimulated and the results of that interrogation are recorded. An unambiguous protocol describes how models are used to run certain simulated experiments, for instance to generate figures or lead to conclusions in scientific papers. This information can be provided by sharing simulation codes; or, again, more reproducibly by providing detailed machine-readable instructions on how to run simulations (such as those provided by SED-ML (Waltemath et al., 2011) or via our existing Web Lab and its protocol descriptions; more on these later).
3. **Model development** — reporting on models at present generally takes the form of their final parameterised equations. Documentation on how models were built (in terms of choosing equations to use, fitting their parameters to experimental data, and validation against other experimental data) is usually insufficient to recreate models accurately, or even missing entirely. Only with this information can you decide whether a model is going to be useful for your setting, or whether you are breaking any assumptions involved in creating it. In this paper we propose that we need a reproducible definition of model development.

**Table 1:**
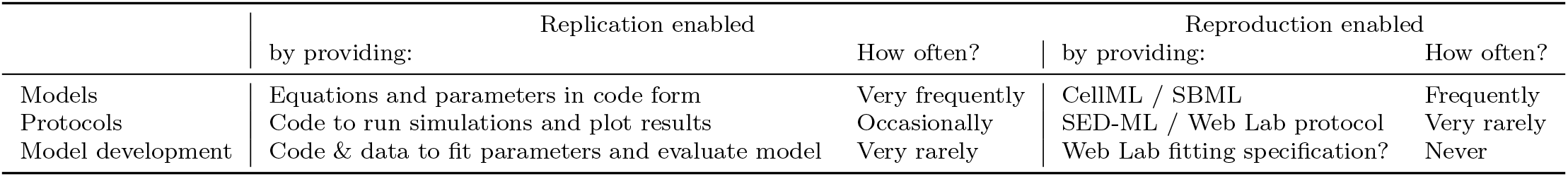
Replicability versus reproducibility in cardiac electrophysiology modelling studies. Here we list what provisions enable replication and reproduction of cardiac electrophysiology modelling studies, and also estimate how often they feature in published studies. Our plan is for the next version of the Web Lab to become the missing tool for making model development reproducible.

The academic cardiac modelling community’s current standard is that replicable models should generally always be provided, at least for action potential models: nearly all studies now provide model code for replication, and most also provide CellML for reproducibility. These efforts are certainly to be applauded and it has taken a large amount of effort, primarily from the CellML team, to get to this point.

Some authors provide their simulation codes to enable replication of protocols and simulated experiments (e.g. Chang et al., 2017), and some journals insist on this; although it remains far from universal in reporting at present. Very few authors provide reproducible definitions of protocols to repeat simulated experiments using their models such as those provided by SED-ML or the Web Lab: this is understandable, as these ‘protocol definitions’ are only just beginning to support all the features that are needed, and tools which support/implement them easily are still being developed.

The development of a full model itself — which involves parameter inference from data, potentially model selection, and then evaluation of model performance — is very rarely even replicable, as this requires complete implementation details for all forward simulations and inverse problem algorithms in addition to providing digitised experimental data linked to particular protocols. The only replicable instance of a full model’s development we could find was Beattie et al. (2018) (used as our case study for reproducible model development below). We hope that readers of this article can point us to more examples. Model development has never been reproducible, partly because it depends on both models and protocols also being reproducible. So the focus of this paper revolves around the steps involved in making model development reproducible, our plans to facilitate this with the Cardiac Electrophysiology Web Lab, and a pilot implementation.

Reproducible reporting standards for model development are especially important for understanding a model’s *provenance.* Many questions are raised when attempting to reproduce the development of a model: what assumptions were made in its construction? Which experiments must be performed to parameterise the model (or re-parameterise for new conditions, e.g. cell types/species/temperatures)? How should experimental data be post-processed before being used to fit parameter values? How much information for constraining the parameter set do the model builders consider each experiment to contain (Fink and Noble, 2009) — relatedly, what fitting/inference algorithms or heuristic approaches used which datasets to generate which parameter values? How was the parameterised model tested/validated, using which experimental protocol and datasets? At present these aspects can be reported very poorly, if at all.

This lack of reproducible documentation on model development makes models increasingly static, as they are difficult to update in response to newly available data, or more advanced understanding of sub-cellular processes, and one may introduce errors or lose desirable behaviour as one attempts to adjust them. Ill-defined model provenance can also make it difficult for modellers or experimentalists to choose which model to adopt from the myriad formulations that have been proposed to explain similar cardiac phenomena.

Knowing the provenance of models is especially important in the field of cardiac models, which are often chimeric, employing data from multiple sources due to the difficulty of gathering data sufficient to constrain today’s complex models from a single source. Cardiac modellers often borrow parameters or entire subunits from other models that may have been constructed for different systems or under differing experimental conditions (Cannon and D’Alessandro, 2006; Niederer et al., 2009). While efforts are taken to adjust model components for differences in species and/or experimental conditions, as well as to maintain certain macroscopic properties, manual tuning practices are unlikely to locate truly optimal parameters (Krogh-Madsen et al., 2016).

When there exist a large number of parameterisations that describe the data equally well, the model is deemed to be *unidentifiable*, and the modeller may wish to consider either an alternative simpler model formulation, alterations to experiments to provide more information on parameters, or both (Raue et al., 2011). Considering overly-simplistic objective functions in parameter tuning, such as matching just biomarkers from a single action potential, may lead to unidentified parameters that can cause models to yield unexpected and erroneous behaviour when tested under new contexts of use (Daly et al., 2017). While model identifiability is a recognised problem in cardiac cell models, its assessment has yet to be adopted as a standard practice during modelling studies, perhaps due in part to the competing methodologies for doing so (Milescu et al., 2005; Fink and Noble, 2009; Csercsik et al., 2012; Sher et al., 2013; Daly et al., 2015, 2017). Unidenti-fiability of parameters may explain some cases of ostensibly similar models yielding qualitatively differing predictions under certain protocols (Cherry and Fenton, 2007; ten Tusscher et al., 2006; Niederer et al., 2009; Fink et al., 2011; Cooper et al., 2016).

Finally, reproducible reporting standards are important for understanding the effects of uncertainty and variability in biological models. Biological data is invariably affected by many sources of variation, such as measurement error, intrinsic variation (e.g. “beat-to-beat” variability between recordings on a single cardiomyocyte), and extrinsic variation (variation between individuals in a sample, e.g. inter-cell variability in a sample of cells, or inter-individual variability in a population). This leads to uncertainty about (or variation in) the optimal model parameters to describe the experimental data, and characterisation and interpretation of this uncertainty can give us insights into biological variation or even the suitability of a given model to explain data (Mirams et al., 2016). Cases of extreme variation in optimal model parameters may indicate the unsuitability of the model to represent the system, as it reduces faith in a direct biological interpretation of each parameter.

In previous work, we sought to address the first two kinds of reproducibility listed above in cardiac modelling studies through the development of the Cardiac Electrophysiology Web Lab. The Web Lab is an online resource for the specification, execution, comparison, and sharing of simulation experiments and their results (Cooper et al., 2016). This Web Lab was built using the functional curation paradigm (Cooper et al., 2011): models are specified using a standard format such as CellML, protocols are specified in a custom language capable of expressing a vast range of experimental procedures, and interactions between the two are mediated by a domain-specific ontology. This allows for modular interactions between models and protocols, allowing for multiple models to be compared under a common protocol or vice-versa, extending the capabilities of contemporary online tools for analysing/comparing cellular model predictions such as WholeCellSimDB (Karr et al., 2012, 2014). The Web Lab additionally provides visualisation tools to aid in these comparative studies, as well as visibility settings that allow models and protocols to be developed in private before being shared with the community.

In this paper, we will discuss our plans for, and initial progress towards, integrating experimental data and reproducible model development into the Cardiac Electrophysiology Web Lab. We will show how the addition of experimental data, annotated and linked to an experimental protocol, can facilitate model/data comparison, model selection, rigorous documentation of model provenance and development, and even automated model reparameterisation and identifiability analysis. In section 2 we describe the steps needed to create such a tool (which we will refer to as *WL2*) from our original implementation (which we call *WL1*). Section 3 showcases a prototype implementation of WL2, with which we reproduce the results of a Bayesian modelling study of the hERG-encoded ion channel conducted by Beattie et al. (2018), shown in section 4. This study of a contemporary cardiac model serves as a proof-of-concept of our WL2 design, and demonstrates its ability to facilitate the fitting of models to data and to provide information about the uncertainty in the obtained parameter values. Finally, we discuss the remaining challenges for a full implementation of the WL2 design as well as the opportunities it creates.

## 2. Road map

We now outline the steps required to establish an improved Web Lab, WL2. An overview of each step and the new capabilities it facilitates is shown in Table 2.

**Table 2:** An overview of the steps needed to go from WL1 to WL2, and the capabilities added at each step.

|  |  |  |
| --- | --- | --- |
| Step 1 | Adding annotated data | Comparing arbitrary data sets<br>Structured queries |
| Step 2 | Linking data to protocols | Comparing experimental protocol results<br>Documenting data provenance |
| Step 3 | Comparing data to predictions | Checking model applicability<br>Documenting model provenance<br>Continuous testing of models |
| Step 4 | Fitting models to data | Driving model development<br>Documenting model development<br>Quantifying experimental variability<br>Identifiability checking<br>Protocol design<br>Validation |

### 2.1 Step l: Adding experimental data

WL2 will introduce the capability to upload, store, and display experimental data. This will open up a new range of uses for the Web Lab such as comparing published data sets or checking new experimental data against gold standard reference data. A crucial step here is that data should be *annotated* with information about its origins, e.g. species, cell-type, experimental conditions. In part, these annotations should be free-form, allowing experimenters to detail the specifics of their data. However, including *structured annotations*, for example following the MICEE standard proposed by Quinn et al. (2011), will allow Web Lab users to perform structured *queries* on the data set. Such a well annotated, searchable data set, would make it easy to compare all data from a specific species, to compare experiments at different temperatures, or to investigate biological variability between data sets with identical experimental conditions. It is our aim that inclusion of these features will make the Web Lab an invaluable “first-stop” community-wide resource enabling, for example, experimenters to check their results against the literature, analysts to compare drug-block data in different cell-types, or for researchers to find data on the effects of genetic mutations.

Electrophysiological measurements are typically contaminated by sources of error such as leak, drift, noise, or capacitance artefacts. The precise nature of these errors — and therefore the best way to compensate for them — is dependent on the measurement method used, rather than the underlying physiology. Therefore, we do not at this time propose to automate pre-processing operations such as filtering or leak subtraction at this time. Instead, our ideal data source would allow storage of data in three parts: (1) a raw unprocessed file, representing the quantity of interest plus various sources of error, (2) a pre-processed file, in which any experiment-specific pre-processing has been performed, and (3) code to reproduce the pre-processing process, to be run off-line. This set-up would allow pre-processing code to be inspected, reviewed, and re-used. It is also worth noting that some types of filtering (e.g. fitting a straight line through noisy data) presuppose a certain structure in the data, and so mix modelling with pre-processing. By having the raw data available online such work could be accommodated.

### 2.2 Step 2: Linking data to experimental protocols

A crucial step in using data sets on WL2 will be to link them to an *experimental protocol*. For example, for ion current experiments this would include data such as the temperature and chemical composition of the pipette and bath solutions, and also the complete voltage-step protocol. Similarly, for cell-level data such as an action potential duration (APD) restitution curve, this would consist of a series of cycle-lengths to test and a description of the post-processing steps required to measure the APD.

In Cooper et al. (2011), we presented a formalism to encode the experimental conditions, procedure, and post-processing operations — the sum of which we termed the *protocol* — in a machine-readable format that can be used to run simulations. This language defines six foundational operations on n-dimensional arrays (such as time-series measurements of e.g. voltage, current, or ionic concentrations) from which more complex operations (e.g. peak current detection, APD measurement) can be formed. For examples, please see the current WL1 implementation at https://chaste.cs.ox.ac.uk/WebLab.

An important feature of the WL1 protocol language is that it is model independent; protocols it describes can be used to run simulated experiments with any suitably annotated model. We demonstrated the use of this language for functional model comparison and debugging in Cooper et al. (2016). By linking these protocols to experimental data sets, WL2 can become a platform for rigorous documentation of experimental data, with a full description of the protocol and post-processing steps required to re-obtain that data. This will also facilitate a more careful comparison of different data sets describing the outcomes of similar protocols.

### 2.3 Step 3: Comparing model predictions to experiments

Once annotated experiments are available, along with machine-runnable descriptions of the experimental protocol and a database of annotated models, it becomes possible to compare experiments to model predictions (Cooper et al., 2015b). This has several important applications.

#### Checking model applicability

Many investigations into cardiac cellular electrophysiology start with the selection of a suitable computational model. Yet until the publication of WL1 there has been no easy way to look up and compare basic model properties (i.e. what outputs the different models predict given a certain protocol). WL1 introduced this ability to compare model predictions systematically, and with WL2 it will become possible to validate model outputs against a variety of data sets. Users wishing to investigate a specific aspect of cellular electrophysiology, for example calcium handling, could start by selecting a series of virtual experiments relating to the phenomena of interest, and then compare models specifically on how well their predictions match the data in these respects.

#### Documenting model provenance

With this new set-up, it becomes possible to describe a model’s provenance in a systematic way, by making formal statements such as ‘parameters *a, b*, and *c* in model m derive from data set *d*’. If, in addition, model sub-components (e.g. ion-current models or calcium buffering equations) are linked with statements such as ‘model *m*_1_ relates to model *m*_2_ via inheritance of sub-model *I_Na_*’, it will become possible to trace the origins of model equations and parameters. Because this provenance can be extremely complicated (Niederer et al., 2009; Bueno-Orovio et al., 2014), having a shared community record of such relations will be extremely useful for the electrophysiology community.

#### Continuous testing during model development

A consequence of electrophysiology (EP) models’ complicated history is that modellers adapting a specific aspect of a model (say the sodium channel formulation) may not be familiar with details of other parts of the model. In addition, the code and data used to run all experiments that were originally used to validate the model are not usually available. With WL2, it becomes possible to encode all these experiments (or better, to re-use ones already available) and link them to the relevant experimental data set. This creates a large body of test-cases that a model developer can use to test any updated model formulations against, to ensure novel additions do not undo the efforts of previous modellers.

### 2.4 Step 4: Fitting models to experimental data

At this point, the Web Lab will allow users to compare experimental outcomes and model predictions not just qualitatively (i.e. visual inspection), but also *quantitatively*. A next step then, is to let the user define some measures that quantify the model/experiment mismatch (or alternatively the goodness-of-fit), and to introduce algorithms that *fit* models to data by systematically adjusting selected model parameters until the predictions match the observed results.

We distinguish two main types of fitting. In *optimisation*, the mismatch between model prediction and experimental outcome is quantified by some measure of error, and an optimisation algorithm is used to reduce this error to a minimum. The outcome of this process is a single set of ‘best’ parameter values. In *statistical inference*, the difference between the model prediction and the observed data is treated as a random variable (e.g. due to measurement noise), and an inference algorithm is used to quantify the likelihood that different parameter sets gave rise to the observed data. This results in a *distribution* of parameter values, each with an associated likelihood.

Further distinctions can be made depending on how the experimental outcome is defined. First, many analyses start from *summary statistics* of the data, for example a current-voltage relation (IV-curve) when measuring ion currents, or a steady-state APD dose-response curve when measuring the cellular AP under drug action. In this method, a certain amount of data is discarded to simplify the (historically manual) analysis process on the assumption that the remaining data will fully characterise the phenomenon of interest. By contrast, *whole-trace fitting* uses all available data and does not require this assumption. Several publications have pointed out the benefit of the whole-trace method for the analysis of ion-currents (Hafner et al., 1981; Willms et al., 1999; Lee et al., 2006; Fink and Noble, 2009; Buhry et al., 2011; Beattie et al., 2018). Secondly, data from different experiments (e.g. in different cells) can either be averaged before processing, or can be processed on an individual basis. Averaging before processing can lead to distorted results, as shown by e.g. Pathmanathan et al. (2015).

In WL2, an outline of which is shown schematically in Fig. 1, we plan to support all of the modes of fitting listed above. An example using our prototype implementation is given in sections 3 and 4. Supporting model fitting will have several important applications.

**Figure 1:**
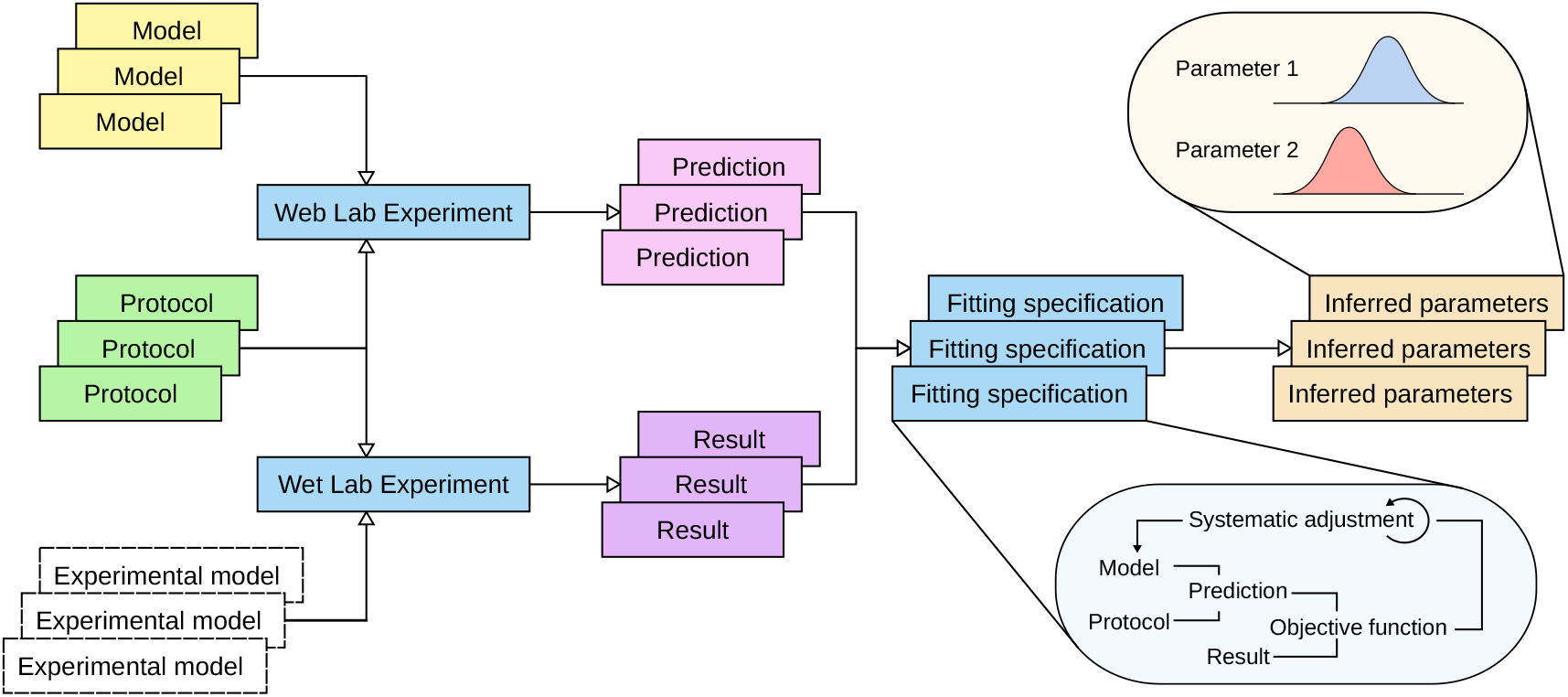
A schematic overview of WL2. Experimental protocols, applied to biological models (e.g. myocytes, expression systems) give rise to experimental results. The association with a protocol, in combination with additional metadata, provides users with a thorough overview of how the data was obtained. Applied to computational models, the same protocols provide predictions. As in WL1, protocols are written in such a manner that they can be applied to several models on the Web Lab, and their predictions can be compared. A new feature will be the ability to compare predictions to predictions, experimental results to results, and results to predictions. By comparing experimental results and predictions from the same protocol, a fitting process can be initiated, leading to a set of parameter values represented either as singular points (optimisation) or distributions (inference).

#### Driving model development

EP models are based on experimental data from a variety of sources, including measurements in different species and under different experimental conditions (Niederer et al., 2009). Once the infrastructure to refit models automatically is in place, it should become a straightforward task to update model parameters whenever new and/or more appropriate data sets are available (see also Box 4 in Cooper et al., 2015b). Similarly, if no model can be reparameterised to fit the data, this is a strong indication that changes to the model equations are required. In this manner, WL2 could be a driving force behind electrophysiological model development.

#### Rigorously documenting model development

With the addition of fitting algorithms, WL2 will allow a large part of model development to be rigorously — and reproducibly — documented. This would dramatically increase the reproducibility of electrophysiology modelling work, and allow the kind of close scrutiny of model-building work that is required if EP models are to be used in safety-critical applications such as drug testing (Mirams et al., 2012) and clinical risk assessment (Hoefen et al., 2012).

#### Quantifying experimental variability

Using statistical inference methods, we can find distributions of model parameter values that provide good fits to experimental data, and quantify the likelihood of each (Daly et al., 2015). This allows us to quantify variability and uncertainty in single parameters, but also to investigate correlations between parameters (Johnstone et al., 2016b). The application of these techniques is a first step towards untangling experimental error from biological variability (Mirams et al., 2016; Marder and Goaillard, 2006).

#### Identifiability checking and protocol design

In addition to analysing experimental data, statistical inference methods can be used as a tool to design experimental protocols. If a broad range or ranges of parameter values are found to be equally likely candidates to fit a data set, this can be a strong indication that the protocol does not provide all the information needed to identify the model’s parameters. Such a result may highlight that the model is fundamentally *unidentifiable*, but it can also highlight the shortcomings of a particular protocol (i.e. it does not trigger all the phenomena the model was intended to capture). By using inference to check if a protocol provides enough information, it becomes possible to test and optimise a protocol, e.g. by removing steps that are found not to add new information (Fink and Noble, 2009; Clerx et al., 2015; Beattie et al., 2018). By making these features widely available, WL2 can aid experimenters in both protocol selection and design.

#### Validation

An important step in developing a biophysical model is *validation*, that is assessing how well your fitted model represents reality (Pathmanathan and Gray, 2013). In our domain this is most easily done by running extra experiments that were not used to fit the model, i.e. a different protocol. It is vitally important that protocols and recorded experimental data for validation are associated with the model development process, and available for display and comparison within WL2. We did not denote this step in Fig. 1 to keep the figure simple, but in full it should feature additional line(s) from the Inferred Parameters back to simulations and experiments with a new protocol. Validation is especially important when extending an existing model. Without a complete description of experimental results that were predicted/recreated with an existing model, it is impossible to know whether a new version of a model retains all the capabilities of the original one.

## 3. Prototype implementation

We now discuss the implementation of a prototype WL2, focused on performing statistical inference over parameters of single-cell EP models and sub-models (e.g. ionic currents), and demonstrate its use by reproducing a result from a hERG modelling study conducted by Beattie et al. (2018). In this study, a novel voltage-step protocol was applied to cells expressing the hERG ion channel, and the recorded current was fitted with a 9-parameter Hodgkin-Huxley model, summarised below in its equivalent Markov-model formulation (for an explanation of the relation between these see Keener and Sneyd, 2009, vol. 1, p150).

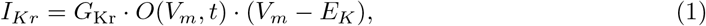

with the conductance parameter *G*_Kr_, and the open probability *O*(*V_m_,t*) given by the system of equations

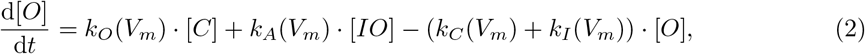

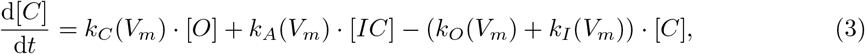

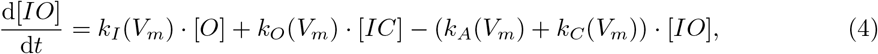

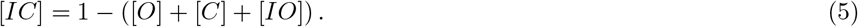

The rates are voltage-dependent functions, each parameterised by two scalar values as follows:

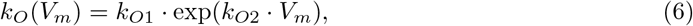

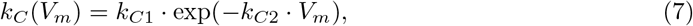

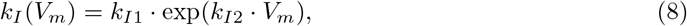

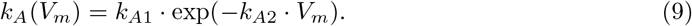

A *Bayesian inference* method was then applied to find the parameter values that provided the best fit. As discussed in section 2.4, this inference method finds not just a single set of parameter values, but the distribution of all likely parameter values, known as the *posterior distribution*.

### 3.1 Simulation Specification

Finding the best parameters, either using optimisation or inference, involves repeated simulations, and so WL2 must provide the user with an interface in which to specify how model simulations should be performed. Our prototype WL2 builds on the design of WL1 (Cooper et al., 2016), and thus adheres to the functional curation paradigm separating the notion of a model (the underlying mathematics representing the system) from the protocol (the manner in which the system is stimulated and observed). As in WL1, this modular design allows for the re-application of experimental protocols to new model formulations, or the comparison of competing model formulations under a given experimental set-up. Users of the WL2 prototype upload separate model and protocol files, represented respectively in CellML and functional curation protocol syntax, in order to specify and share the results of simulations. We have chosen to use the custom protocol language rather than SED-ML due to the latter’s adherence to a “one model, one experiment” paradigm (simulations are explicitly linked to models rather than mediating interactions through a domain-specific ontology), which lacks the modularity of functional curation. Additionally, the functional curation protocol language supports more complex nested simulations and provides greater post-processing power than SED-ML, allowing it greater expressive capability. As SED-ML evolves we hope to use it to specify protocols and simulation algorithms in future versions of the Web Lab, and anticipate our protocol language serving as a test-bed for new features for the community standard.

Model parameters in the CellML files should be tagged with metadata annotations that allow them to be externally read and adjusted by simulation protocols, and thus there exists a notion of *compatibility* between model and protocol, which requires that all variables referenced in the protocol exist as tagged entities in the model. The exact structure and interpretation of these files are given in Cooper et al. (2011).

In addition to the simulation specification, the WL2 prototype requires two files to complete the specification of a fitting problem: a file containing experimental data, and a *fitting specification* that shows how this data is employed to constrain the model. The content of these files will be discussed in the subsequent sections.

### 3.2 Data Specification

In our prototype WL2, data for fitting experiments is provided in a separate file. Each entry should directly correspond to an output of the simulation protocol,and consist of a series of data arrays with unique names, as the functional curation protocol language possesses a postprocessing library to enable such one-to-one matching (Cooper et al., 2011). Much like the functional curation paradigm, the use of unique names to mediate interactions between the data and the other components of a modelling study allows for experiments to be easily updated when new data become available.

In our prototype implementation, data are supplied in a comma-separated values (CSV) file, where the first row specifies names for each variable, and associated columns specify the corresponding data. We note that this structure currently expects zero- or one-dimensional data for each named variable (although higher-dimensional arrays may be specified in flattened form), as this was sufficient for our test case, but that the exact data representation can easily be changed at a later stage. In the hERG current fitting experiment, the data file contains two columns of equal length representing a series of (time, current) pairs. Our current implementation requires that the units of the data provided match those of the corresponding entities in the simulation protocol, though future iterations of WL2 will allow for the specification of units and handle necessary translations in a manner similar to functional curation.

### 3.3 Fitting Specification

The final component of a parameter fitting experiment is the fitting specification, which makes use of a custom language that we will introduce in this section by means of a working example.

The fitting specification takes the form of a JSON-formatted text file (http://www.json.org). These types of files contain a series of named values, where each value may be either a string of characters, a numerical value, a list of values, or a nested JSON object. We represent the contents of a fitting specification for the hERG model based on the fitting experiment described by Beattie et al. (2018) (with the number of CMA-ES optimisations and number of MCMC iterations reduced for the sake of improving runtime) in Table 3, and will discuss the interpretation of each required entry below.

**Table 3:** Entries in the fitting specification for the hERG ion channel model. The value associated with the “algorithm” entry is a string of characters, and is represented as is, while all other value entries are nested JSON objects, and are presented in “key=value” format for clarity. This is also true for the prior specification, which is represented separately in Table 4 due to its size.

| Fitting specification entity | Value |
| --- | --- |
| algorithm | AdaptiveMCMC |
| arguments | cmaOpt=5, cmaMaxFevals=20000, burn=50000, numIters=100000 |
| output | exp_IKr=IKr |
| input | exp_times=t |
| prior | (see Table 4) |

The first entry in the fitting specification is the “algorithm” to be used for parameter fitting, which is specified by a unique identifier. In the case of the hERG experiment, this value is “AdaptiveMCMC”, which corresponds to the adaptive-covariance MCMC algorithm described by Haario et al. (2001). While we are interested in moving towards more sophisticated means of uniquely specifying algorithms, such as KiSAO IDs (http://co.mbine.org/standards/kisao) in future iterations of the Web Lab, such ontologies do not yet support the full range of Bayesian and approximate Bayesian inference algorithms that we are considering for inclusion in WL2. Once we refine the list of algorithms we support, we will lobby for their inclusion in KiSAO (or another accepted ontology) and adapt to this new form of algorithmic specification in future iterations of the Web Lab.

In our prototype implementation, the adaptive MCMC algorithm uses a Gaussian likelihood function, which is commonly assumed for time-series data. However, the prototype could easily be extended to allow users to specify different likelihood functions, by adding an “objective” entry to the fitting specification language.

The next entry we consider is a dictionary of “arguments,” specific to the chosen fitting algorithm. In the example shown in Table 3, these include the standard arguments for MCMC — the total number of iterations “`numIters`” and the number of iterations discarded as burn-in “`burn`” — as well as two new arguments “`cmaOpt`” and “`cmaMaxFevals`” which deal with the selection of a starting point for MCMC. These arguments tell the back end to first run a series of 5 random restarts of a global optimiser, the “Covariance Matrix Adaptation Evolution Strategy” (Hansen and Ostermeier, 2001) to choose a starting point for the MCMC chain. In the final version of WL2, a full list of available named algorithms, along with details of their operation and adjustable parameters, will be made available on the web site.

The next two sections, “`input`” and “`output`”, deal with matching experimental and simulated data. The “`input`” section details a list of named inputs to the simulation protocol (“`exp_times`”, in this instance) which are matched to named entries in the data file (“`t`”, in this instance). Here this tells the simulator to generate outputs at the times specified in the data file, and removes the need to alter the functional curation protocol when new data are collected. The “`output`” section tells the objective function which named outputs from the simulation protocol are to be considered (“`IKr`” in this instance), and which named entries in the data file (“`exp_IKr`”) they are to be compared against using the objective function. This allows data files to be used that do not adhere to the same naming conventions as the associated simulation protocol, again avoiding the need to alter simulation protocols when new data are acquired.

Finally, we consider the *prior distribution*; this represents our ideas about the likely values of the parameters before we start the fitting experiment, and is commonly given as a uniform distribution with some lower and upper bound (which typically maps onto the expected maximum physiological range of the parameter). In the WL2 prototype, the prior is specified as a nested JSON object identified by “`prior`” (and represented separately in Table 4). Our implementation currently supports only uniform priors (specified by a two-membered list containing a lower and upper bound) or point distributions (a single fixed value), but more can easily be added. Additional constraints on the model may be implemented as assertions within the simulation protocol itself. This methodology was used to eliminate parameterisations leading to non-physiological values for rate constants (*k_O_*, *k_A_, k_I_*, and *k_C_*), assigning them zero probability in a similar manner to the 2D prior employed by Beattie et al. (2018).

**Table 4:** Prior distribution specified within the fitting specification for the 9-parameter hERG model. This prior is adapted from Beattie et al. (2018), who employ a wider prior in their MCMC inference but define this region as most likely to contain the optimal parameters. Parameters respect the shortened naming conventions of Equations (1)—(9) for clarity. An additional parameter, “obj:std”, controls the observation noise standard deviation, part of the Gaussian likelihood function, and is set to a fixed value in this example (although in general it could be learned too).

| Parameter Name | Prior Range |
| --- | --- |
| $k_{O1}$ | Uniform(1e-7, 0.1) |
| $k_{O2}$ | Uniform(1e-7, 0.1) |
| $k_{C1}$ | Uniform(1e-7, 0.1) |
| $k_{C2}$ | Uniform(1e-7, 0.1) |
| $k_{I1}$ | Uniform(1e-7, 0.1) |
| $k_{I2}$ | Uniform(1e-7, 0.1) |
| $k_{A1}$ | Uniform(1e-7, 0.1) |
| $k_{A2}$ | Uniform(1e-7, 0.1) |
| $G_{Kr}$ | Uniform(0.0612, 0.612) |
| obj:std | 0.00463 |

## 4. Prototype results

We now present the results of our test-case fitting experiment, implemented in a WL2 prototype. As with the WL1 implementation, an experiment may be carried out, and its results viewed, by matching a model to a protocol in the ‘Experiments’ matrix view. Within the prototype WL2, the only change to this set-up is that a ‘protocol’ entry now encompasses a simulation protocol, fitting specification, and data file (as described in Section 3). The files used to represent this fitting experiment are described in Section 3 (particularly Tables 3 and 4). Further details, including links to the relevant online resources, can be found in Daly (2018).

After the execution of a fitting experiment, the first thing that the WL2 prototype allows us to do is to compare the data simulated using our inferred maximum likelihood parameters to the experimental data we employed during fitting. In Fig. 2, we see the results of overlaying data simulated using the maximum posterior density parameters returned by our MCMC inference onto the experimental data used to obtain these fitted parameter values. As the MCMC algorithm returns a sample of parameter sets approximating the true distribution over parameters given data, this visual shows us how well the most likely parameter set captures the observed behaviour. The close agreement between these traces suggests that the inference strategy has produced a distribution over parameterised models that captures observed behaviour well, which mirrors the findings of Beattie et al. (2018). Had we employed an optimisation strategy instead of a Bayesian inference strategy we could also use the maximum likelihood values, which (in this case) would be the same as the least squares optimum, and simply compared the data simulated under these optimal parameters with experimental observations.

**Figure 2:**
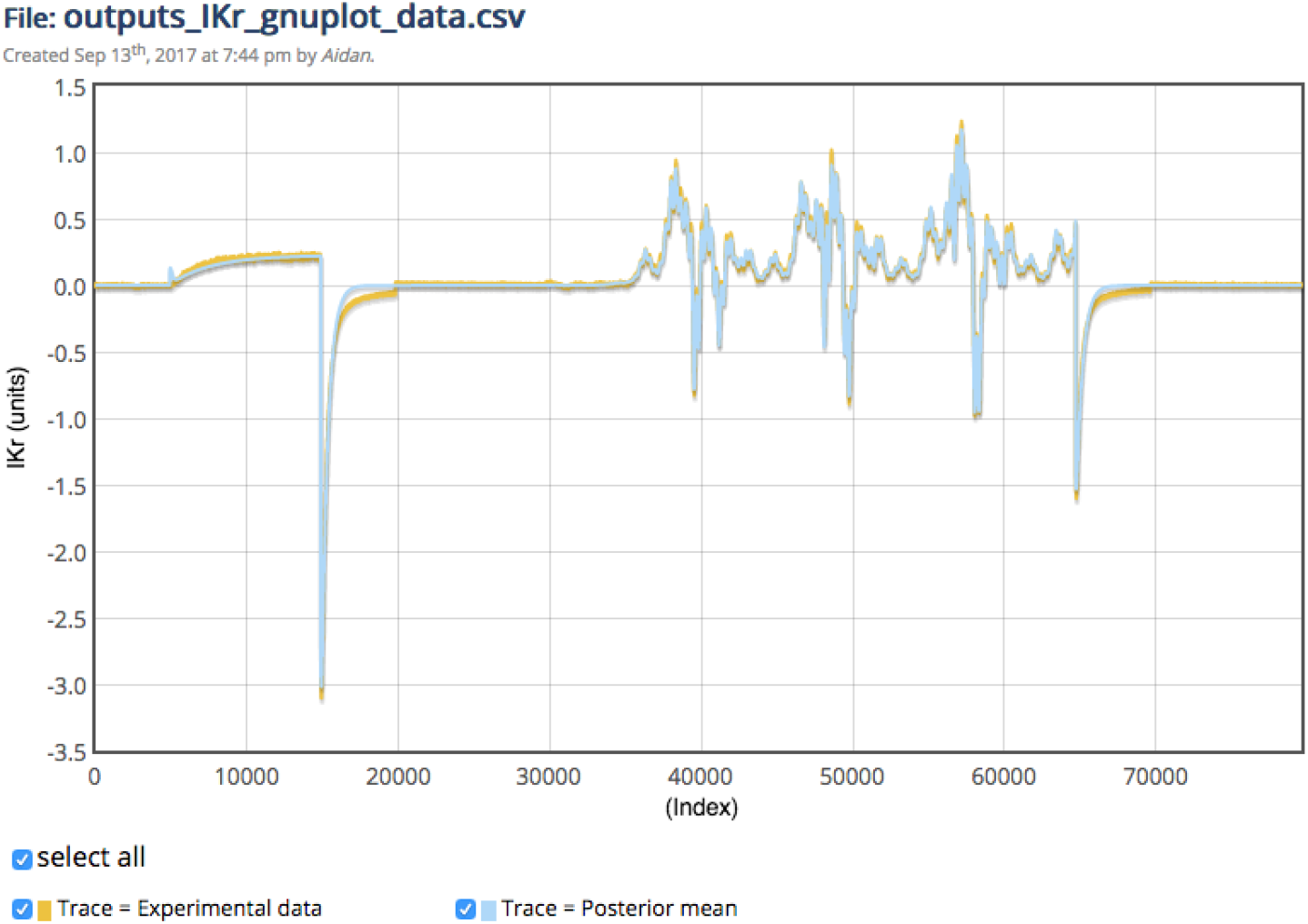
A screenshot of the WL2 prototype: data simulated under maximum posterior density parameters of the hERG model produced by MCMC overlaid with experimental data. Indices on the x-axis correspond to time in seconds with sampling every 0.1ms. The comparable plot in the original model publication is Fig. 4 in Beattie et al. (2018).

The prototype WL2 additionally provides tools for visualising uncertainty about optimal model parameters in inference studies, as characterised by the *marginal distribution* over each parameter in the posterior estimate. In Fig. 3 we see a histogram produced by the prototype WL2 in order to represent variation of the *k*_*A*1_ parameter of the hERG model over the posterior returned by MCMC. We see that this distribution is very well-constrained about a single modal value, which indicates the presence of a unique optimal value for this parameter about which the model is very sensitive to variation (with even small changes in the value of the parameter leading to a large drop-off in likelihood). The posteriors for the remaining model parameters (see both Beattie et al. (2018) and Daly (2018)) show similar constraint. While this could potentially indicate a well-defined and narrow local optimum, the fact that we employed multiple starts to our initial CMA-ES optimisation strengthens our belief that this is indeed a global optimum, and supports our belief that the model is uniquely identifiable under the current experimental set-up. Had one or more parameters shown multi-modality or flatness in their marginal posterior distribution, we would conclude that the data did not provide enough information to uniquely constrain all model parameters (Siekmann et al., 2012). In this manner, examination of the model using the WL2 prototype may reveal unidentifiabilities in the model, which may require an alternate model formulation or experimental protocol to address depending on the character of the variation (Daly et al., 2017).

**Figure 3:**
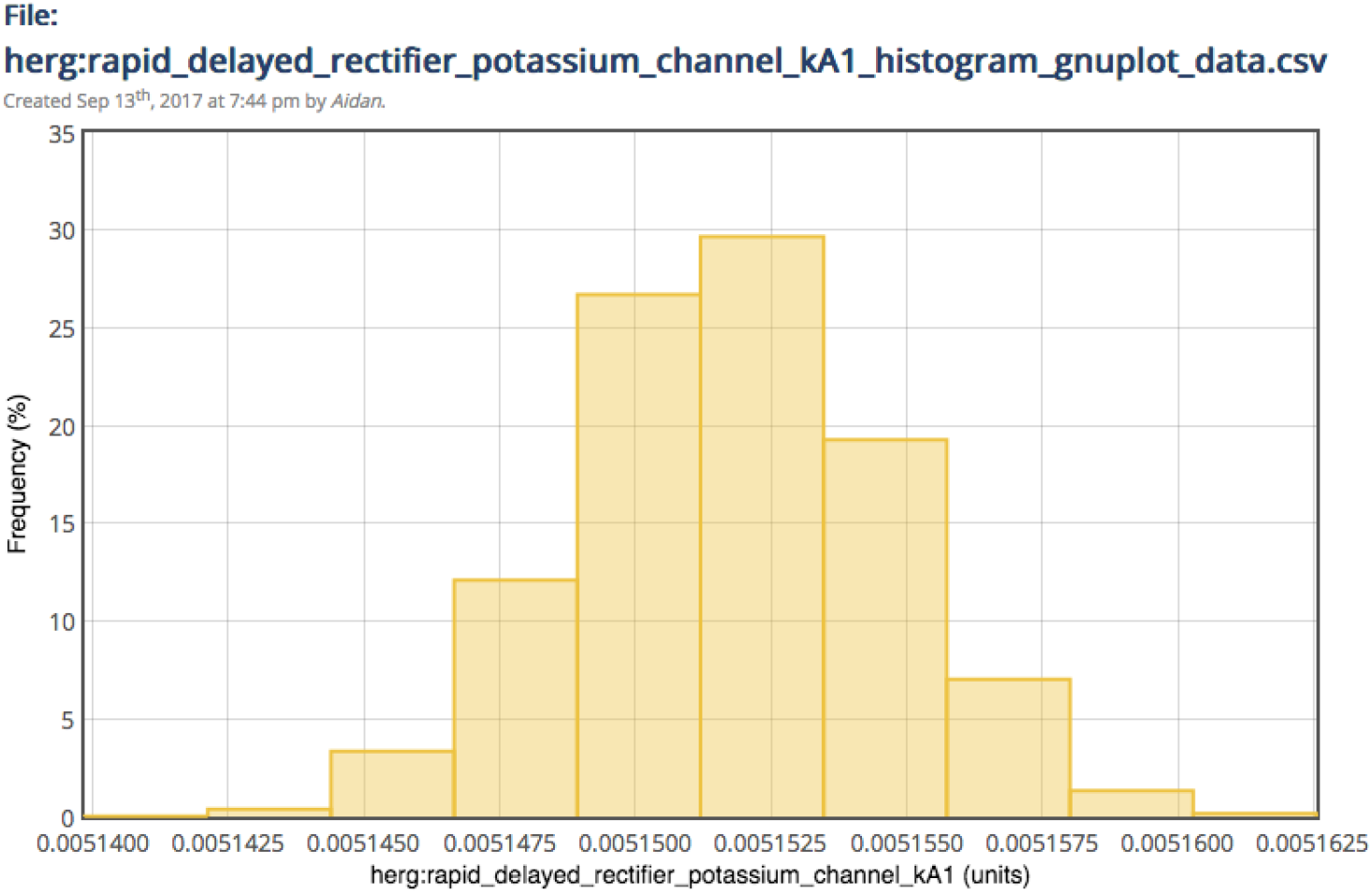
A screenshot of the WL2 prototype: visualisation of marginal variation over kinetic rate parameter *kAi* of the hERG model in the posterior distribution returned by MCMC. Comparable histograms in the original model publication are shown in Fig. 4 of Beattie et al. (2018).

## 5. Discussion

We have laid out the steps required to extend the Cardiac Electrophysiology Web Lab with experimental data, and discussed the advantages this will bring. With a prototype implementation of this WL2 we have shown the feasibility of using the Web Lab to perform statistical inference, the most technically challenging of the features discussed in our road map. We now discuss remaining challenges for a full implementation of WL2 and its adoption by the electrophysiological community, as well as the opportunities some of these challenges present.

### Providing incentives for model annotation

The option to annotate models, sub-models, and variables has long been present in model-sharing languages such as CellML (Hedley et al., 2001), but is not widely used (see e.g. https://models.cellml.org/electrophysiology). Similarly, initiatives such as the MICEE Portal (https://www.micee.org/), which is intended to store annotations for published experimental data, have so far failed to garner widespread adoption. Clxearly, there has not been sufficient incentive for scientists to provide annotated models and/or data sets. However, very few tools *make use* of model or data annotations, so that the updated Web Lab’s annotation use could serve as a prime example of the benefits of annotation. As soon as a link is established between experimental data and the models that use it (section 2.2), and refitting (parts of) models becomes a routine task, questions of model-data provenance will arise more naturally, and the need for well-annotated data will be felt by a wider audience.

### Creating a community repository for electrophysiological data

A second challenge related to the addition of experimental data to the Web Lab is dealing with the administrative and financial burdens of storing and providing access to valuable data for an indefinite period. Conflicts of interest might best be avoided if this responsibility was placed in the hands of some independent multi-centre organisation, and so it is clear that this task should be undertaken separately from establishing WL2. One of the many opportunities the establishment of such a repository would bring, is that it would make it easier for multiple groups to tackle the same problem, using the same data set. For example, the *PhysioNet challenge* is an annual competitive event where computational biology groups around the world are challenged to provide the best analysis of a particular biophysical signal (usually an ECG) from the PhysioNet/PhysioBank repository (Goldberger et al., 2000). If a repository for cell electrophysiological data were to be established, similar events could be run to tackle questions in the cell electrophysiology domain. One possibility is that such a repository might be linked to a community-based journal, and responsibility would be assumed by the journal publisher. It would also allow the cardiac modelling community to share their wealth of already acquired data for future model development and model verification or validation steps. Preliminary discussions have suggested that leading academic publishers would be open to such an idea, as it is in line with their move towards becoming information platform providers.

### Comparing complex data sets

Biological systems are irreducibly complex; even when only a single ionic current is measured, the ‘background’ is a living cell in which thousands of dynamical mechanisms interact to create and maintain homoeostasis. As a result, any two independent investigations into the same phenomenon will almost invariably differ in some details, some of which may later turn out to be important. For annotation, this means that even when using a standard such as the MICEE draft — which specifies around 100 properties to be recorded as ‘minimum information’ — some details will go unrecorded. It also means that the question of whether two data sets describe ‘the same’ experiment is not always easy to decide. Conversely, many experiments to measure a certain property, for example inactivation of the hERG current, use different voltage protocols (and so are demonstrably not identical) but provide essentially the same information. By enabling data-to-data, data-to-model, and model-to-model comparison, all with excellent support for annotation, WL2 can help bring these issues into much sharper focus.

### Establishing community ontologies

Structured annotation of models, protocols, and data, requires *ontologies*: formal naming systems that allow model variables, protocol properties, and experimental conditions to be uniquely defined. For WL1, we created a custom ontology that defined several model variables, e.g. *membrane-rapid-delayed-rectifier-potassium-current* denotes the current known as I_Kr_, carried by the channel encoded by hERG. In the long-term, this should be replaced by a community agreed-upon ontology, and kept up to date with new scientific discoveries. As with annotation, widespread use of WL2 could be a powerful driving force behind such efforts.

### Integrating with other community tools

Ultimately, we want WL2 to be part of a wider web of community tools and data resources, sharing information via agreed-upon ontologies as discussed above. For this purpose, WL2 will provide not only a user-friendly ‘human’ interface, but also an application programming interface (API). Such an interface (based on the Representational State Transfer, or REST architecture) was already used internally for WL1, but will need to be documented and made publicly available in WL2. We look forward to working with the community to establish the best interface for WL2 (and other tools) to present. A second difficulty relating to interaction, is that not all formats used in WL2 were created with annotation in mind. For example, there is no widely agreed upon method of annotating parts of CSV or proprietary binary data files, and similarly the text-file formats the WL2 prototype uses to specify fitting experiments cannot easily be annotated. However, we believe that, starting from an imperfect implementation, WL2 can help bring clarity and urgency to efforts to establish consensus on such topics. Finally, all entities used in WL2 should have a unique identifier, e.g. a digital object identifier (DOI). As we envision most WL entities (models, data sets) being hosted primarily outside the WL, this not something we can ourselves directly address. However, if simulation protocols and fitting specifications are to be shared with other tools, a solution to obtain such unique identifiers will need to be found.

### Model-agnostic fitting

In WL1, we defined an ontology for well-known variables such as named ionic currents (e.g. the *fast sodium current*), maximum current densities (e.g. the maximum conductance for the fast sodium current), and reversal potentials (e.g. the reversal potential relevant for the fast sodium current). This allows experimental protocols to be written in a model-agnostic manner, that work regardless of the variable names used in the model code.

For fitting specifications to work in a similarly model-agnostic manner, we need to indicate which parameter values should be modified to perform the fit. Whilst some parameters, such as ion current densities, may exist and represent the same quantity in different models, this is not always the case. Model-specific parameters can arise in a number of circumstances, for example when a new biophysical mechanism is postulated, or when a model introduces a deliberate simplification (e.g. Hodgkin-Huxley rather than full Markov models for channel gating). So instead of defining unique names for such parameters, we propose to identify them via their *relationship* to named ontology variables. For example, parameters for the fast sodium current — which is a variable named in our ontology — could be tagged with the property “is a parameter for the fast sodium current”. Further properties could provide a more detailed description, e.g. “influence fast sodium current activation”, or “appears in a reaction rate of the form *ae^bV^*”. A fitting algorithm for current “*x*” could then gather all variables tagged as “parameter for *x*” and vary them in an optimisation, perhaps guided by further properties to add boundaries, parameter transformations, or other “tweaks”.

### Optimisation and statistical inference

A huge number of algorithms exist for optimisation and statistical inference. However, simulators for cell and ion current models usually do not provide information about derivatives with respect to model parameters, which limits the number of applicable methods. Despite recent work by e.g. Syed et al. (2005); Loewe et al. (2016); Moreno et al. (2016); Johnstone et al. (2016a), the best methods to use remain unclear. In addition, setting up a fitting experiment requires detailed knowledge of cellular electrophysiology, so at present developers of statistical methods are unlikely to test their methods on such cases. WL2 provides an excellent opportunity to address both issues. Firstly, by defining a shared interface for optimisation/inference problems we can set up a system where model developers can test out different methods without making changes to their code (for an early version of such a shared interface, see https://github.com/pints-team/pints). Secondly, by working more closely with the statistical community, and using WL2 to make fitting experiments using a standard interface available online, it becomes possible to use published electrophysiology experiments as a test-bed for new algorithmic work. This will result in a mutually beneficial situation for electrophysiology modellers and statistical algorithm designers.

### 5.1 Conclusion

At present developing cardiac electrophysiology models is “something of an art” (anonymous senior cardiac modeller). We would like to see it become a science, defined by an unambiguous algorithmic procedure. Our hope is that in the future a resource such as the Web Lab will provide researchers with everything they need to know to reproduce a model’s development. The Web Lab will list what experimental protocols need to be performed in a wet lab in order to parameterise a model, and then receive data from these experiments and use them to produce parameterised mathematical models. Further in the future we aim to automate the process of developing new models. By including protocol design, optimised experiments could be generated to optimally constrain each parameter, and these could be suggested by the Web Lab in response to the results of previous experiments. Design of protocols for the process of model selection (choosing the most appropriate set of equations and number of parameters to fit) could also be automated, something that is certainly possible for relatively well-defined model structures such as ion channel models. In the future we envisage this protocol optimisation occurring in real time in response to experimental results being recorded, so that multiple rounds of model refinement and new experiments can be performed in one sitting.

In this article we have described our work to date in developing a community-based resource to support the cardiac electrophysiology research community. Our goal is that this resource should become a repository for all aspects of the research of this community: experimental data and protocols, the computational models that are derived from that data and aid in its physiological interpretation, and the instantiations in software of statistical inference techniques that are used to derive those computational models from the experimental data and protocols. Whilst our work as described here focuses on cardiac electrophysiology, the need that we address and the approach that we have used are applicable across a large swathe of scientific research endeavour. In order to make our work as widely accessible as possible, and in the hope that this approach might be adopted more widely in other research domains, all of our work is freely available as open source code at https://github.com/ModellingWebLab under a BSD 3-clause license. Code for the WL2 prototype can be found there in the `fcweb` repository, under the `cardiac-fitting` branch.

## Acknowledgements

A.C.D. was funded by the Rhodes Trust, UK. M.C., J.C., G.R.M. and D.J.G. acknowledge support from BBSRC [grant BB/P010008/1]. K.A.B was supported by the EPSRC [grant numbers EP/I017909/1 and EP/K503769/1]. This work was supported by the Wellcome Trust [grant number 101222/Z/13/Z]: G.R.M. gratefully acknowledges support from a Sir Henry Dale Fellowship jointly funded by the Wellcome Trust and the Royal Society.

